# Warmer temperature accelerates senescence by modifying the aging-dependent changes in the mosquito transcriptome, altering immunity, metabolism, and DNA repair

**DOI:** 10.1101/2025.10.27.684792

**Authors:** Lindsay E. Martin, Jordyn S. Barr, Jean-Philippe Cartailler, Shristi Shrestha, Tania Y. Estévez-Lao, Julián F. Hillyer

## Abstract

**Background:** Global environmental temperatures are rising, which is increasing the body temperature of mosquitoes. This increase in body temperature is accelerating senescence, thereby weakening immune responses and reproductive processes, and shortening lifespan. To determine how warmer temperature and aging, individually and interactively, shape the transcriptome of the African malaria mosquito, *Anopheles gambiae*, we conducted RNA-sequencing and network-analysis in naive and immune-induced mosquitoes that had been reared at 27C, 30C or 32C and were 1, 5, 10 or 15 days into adulthood. Results: We demonstrate that immune induction, warmer temperature, and aging alter the transcriptome. Notably, the transcriptome of 1-day-old mosquitoes is pronouncedly different from older mosquitoes, and importantly, warmer temperature modifies the aging-dependent changes to accelerate senescence. For example, warmer temperature amplifies the aging-dependent decrease in immune gene expression but dampens both the aging-dependent decrease in metabolic gene expression and the aging-dependent increase in DNA repair gene expression.

**Conclusions:** Altogether, warmer temperature accelerates senescence, shaping the transcriptome in ways that alter the mosquitos ability to fight infection and survive in a warming environment.

## Background

Most insects are ectotherms and poikilotherms; they do not use metabolism to regulate their body temperature, and their body temperature fluctuates with the temperature of the environment. As the planet becomes warmer, the body temperature of insects is increasing, which is affecting physiology. At the organismal level, warmer temperature accelerates development, increases metabolism, and alters body composition, but it also reduces body size, decreases fecundity, weakens immunity, and lowers survival (1–15). This is reflected at the transcriptomic level, where changes in temperature modify the expression of genes involved in metabolism, detoxification, and immunity (16–18). Given that mosquitoes transmit disease-causing pathogens, continued temperature-dependent changes in mosquito physiology and survival will have profound consequences on the global burden of disease (14, 19–21).

Aging is another factor that affects physiology. As mosquitoes age, their body deteriorates via a process called senescence (22–24). At the organismal level, aging decreases the mosquito’s protein content, decreases reproduction, reduces immune prowess, and lowers survival (1, 25–34). This is reflected at the transcriptomic level, where aging increases the expression of genes involved in detoxification while decreasing the expression of genes involved in metabolism, autophagy, salivary gland protein production, and the maintenance of cuticular structure (35–38). Moreover, aging modulates the expression of pathogen recognition molecules and antimicrobial effectors (38–41). Although mosquitoes must outlive the extrinsic incubation period of blood-borne pathogens for disease transmission to occur, as mosquitoes get older, their ability to transmit pathogens diminishes (22, 28).

Warmer temperature and aging interact to shape mosquito physiology. Specifically, warmer temperature accelerates the aging-dependent decline in survival (9). Moreover, warmer temperature also quickens the aging-dependent decrease in protein content and the ability to both reproduce and respond to an infection (1, 9, 12, 42–46). We hypothesize that the warming-based acceleration of senescence is reflected at the transcriptomic level, and the examination of transcriptomic changes will shed light on additional senescence-based consequences of warming environmental temperatures.

Here, we elucidate how warmer temperature and aging, individually and interactively, shape the transcriptome of naïve and immune-induced female *Anopheles gambiae*. Using an RNA-sequencing and network-analysis approach, we demonstrate that immune induction, warmer temperature and aging dramatically alter the transcriptome and that warmer temperature modifies the effects of aging. For example, warmer temperature amplifies the aging-dependent changes in immune gene expression but dampens the aging-dependent changes in metabolic and DNA repair gene expression. These transcriptomic changes explain differences in a mosquito’s ability to respond to infection and survive in its warming environment.

## Methods

### Mosquito rearing and experimental overview

A colony of *Anopheles gambiae* (Giles *sensu stricto*, G3 strain; Diptera: Culicidae) was maintained at the standard rearing temperature of 27°C, 75% relative humidity, and a 12h:12h, light:dark photoperiod, as described (9, 12). Eggs produced by this colony were transferred to three environmental chambers with similar humidity and light conditions, but kept at three different temperatures: 27°C, 30°C or 32°C. These temperatures were selected because they simulate higher environmental temperatures that mosquitoes may experience in nature as the climate warms (47, 48). Eggs were hatched in water, larvae were fed koi food mixed with baker’s yeast (2.8:1 koi:yeast ratio), pupae were separated daily, and adults were fed 10% sucrose *ad libitum*.

At each temperature, adult females that were 1, 5, 10, or 15 days after emergence were allocated into two immune treatment groups: (i) naïve (not injected), and (ii) immune-induced by injecting heat-killed *Escherichia coli*. These ages were selected to encompass the physiological changes that occur throughout adulthood (9, 25), and to include ages relevant for parasite development in the mosquito (49). *E. coli* was selected as the immune inducer so that the transcriptomic findings could be compared to prior phenotypic studies that assessed how warmer temperature modifies aging (1, 9, 12, 42). Bacteria were heat-killed to avoid the confounding variable of differing bacterial growth rates at each temperature (50). Because the transcriptional response to heat-killed bacteria wanes over time (51), we selected a 6 h post immune treatment time point and a dose of OD_600_ = 2 to maximally capture transcriptomic changes. The choice of timepoint and dose was validated in preliminary RT-qPCR experiments (Additional Files 1-2: Figure S1, File S1).

GFP-expressing and tetracycline-resistant DH5α *E. coli* were grown overnight in 5 mL of Luria-Bertani (LB) broth in a 37°C shaking incubator (New Brunswick Scientific, Edison, NJ, USA). Prior to injection, bacteria were diluted to OD_600_ = 2, heat-killed at 95°C for 10 min, and cooled on ice for 5 min. Using LB + tetracycline agar plates, heat-killed bacteria were plated to confirm death, and live bacteria were plated to determine the number of colony forming units (CFUs) that would have been in each injection had they not been heat-killed (∼9,300 CFUs per heat-killed injection). For immune induction, 69 nL of heat-killed *E. coli* were injected into the hemocoel via the thoracic anepisternal cleft using a Nanoject III Programmable Nanoliter Injector (Drummond Scientific Company, Broomall, PA, USA).

After immune treatment, naïve and immune-induced mosquitoes were returned to their respective temperatures and provided 10% sucrose *ad libitum*. At 6 h post immune treatment, mosquitoes were briefly anesthetized on ice, groups of 12-15 females were homogenized in 200 µL of TRIzol Reagent (Invitrogen, Carlsbad, CA, USA), and samples were stored at -80°C until RNA extraction.

Five independent biological trials were completed for each temperature-age-immune treatment combination, resulting in 120 samples derived from 1772 mosquitoes. All immune treatments were performed in the morning and all sample collections were performed in the afternoon to avoid possible confounding effects related to circadian rhythms. All graphs were generated using R (version 4.1.0) and assembled using Adobe Illustrator.

### RNA extraction, library preparation, and RNA-sequencing

RNA was extracted using the TRIzol-chloroform phase separation protocol per manufacturer’s instructions (Fisher Scientific, Waltham, MA, USA), and resuspended in Buffer RLT (Qiagen, Hilden, Germany) mixed with 1% 2-mercaptoethanol (Fisher Scientific). RNA was then purified using the RNeasy Mini Kit (Qiagen), treated with DNase on the column (Promega, Madison, WI, USA), and eluted in 32 µL of RNase-free water. RNA concentration and integrity was determined using the Agilent 2100 Bioanalyzer and High Sensitivity RNA TapeStation (Agilent Technologies, Santa Clara, CA, USA).

Library preparation for sequencing was completed using the stranded mRNA NEBNext Poly(A) mRNA library preparation kit with an input of 300 ng of RNA (New England BioLabs, Ipswich, MA, USA). Library quality control was conducted using fluorometric Qubit (Invitrogen) and the 2100 BioAnalyzer. Libraries were sequenced on the Illumina NovaSeq 6000 (paired-end, 150 base pair read) at Vanderbilt University Medical Center’s VANTAGE core facility. The RNAseq data, including raw and normalized gene counts for each sample, and the metadata, including sample information and protocols, were deposited in NCBI’s Gene Expression Omnibus (GEO) database (52), and are accessible through GEO Series accession number GSE284722.

### RNAseq data processing

Sequencing reads were adapter-trimmed and quality-filtered using Trimgalore v0.6.7 (53). An alignment reference was generated from the *A. gambiae* genome (AgamP4.14, VectorBase-66), and trimmed reads were aligned to the genome and counted using Spliced Transcripts Alignment to a Reference (STAR) v2.7.9a with the “– quantMode GeneCounts” parameter (54).

About 20 million uniquely mapped reads were acquired for each sample, resulting in a mapping rate of ∼85%, which is above what is considered sufficient sequencing coverage (55). Sample-level quality control, low count filtering, and normalization to sequencing depth were conducted using the DESeq2 package v1.36.0 (56). Genes counted fewer than five times in three or more samples were removed (57). To investigate the similarities and differences between samples, we first performed Principal Component Analysis (PCA) and evaluated how the samples clustered by immune treatment, temperature, and age. Five samples were removed from downstream analyses because PCA deemed that four samples were outliers (two naïve samples clustered with 58 immune-induced samples, and two immune-induced samples clustered with 58 naïve samples) and one sample was a statistical outlier by Cook’s distance (58). Of the 120 samples, 115 were used in downstream analyses.

### RNAseq sample clustering and Weighted Gene Co-expression Network Analysis

We next conducted iterative weighted gene co-expression network analysis (iterative WGCNA) to (i) identify the biological processes that were most affected by each trait of immune treatment, temperature, and age and (ii) organize the transcriptome into “modules” of genes that may have similar co-expression patterns (59, 60). To do so, we first identified how genes correlate in a network and performed power analysis of the signed network of correlations using the “beta” power selection for iWGCNA’s soft thresholding to reach scale-free topology (61), resulting in a power of 20 for downstream analysis (Additional File 3: File S2). Then, we used module-trait-correlation analysis to identify how each module’s eigengene (the first principal component, akin to a summary measure of the module’s overall activity) correlates with each immune status, each temperature, and each age. We plotted these correlations and their p-values in a heatmap, in which modules with more similar expression patterns are clustered together on the vertical axis (unsupervised hierarchical clustering based on Euclidian distances). We then identified the primary function of genes within a module by over-representation analysis (ORA) with the clusterProfiler v4.2.2 R package (62) that used gene description and gene ontology (GO) terms specific to *A. gambiae* derived from the Vector Base *A. gambiae* PEST assembly GO annotations, version 66.

### Main effect analysis: Immune induction, warmer temperature, and aging

To identify the individual effects of immune induction, warmer temperature, or aging, we next performed differential gene expression (DE) analysis using DESeq2. Genes were considered differentially expressed when they were 2-fold different (log_2_ fold change ≥ 1 or ≤ -1) and had Benjamini-Hochberg adjusted *p* < 0.05 (63). This threshold captures biologically meaningful changes in gene expression (56).

To identify the main effect of immune induction, we conducted a pairwise DE comparison between immune-induced mosquitoes and naïve mosquitoes, irrespective of rearing temperature or age at immune treatment, using naïve mosquitoes as the baseline. For temperature and age, data on naïve and immune-induced mosquitoes were first separated so they could be analyzed independently. To identify the main effect of temperature for each immune treatment, irrespective of age, we conducted a pairwise DE comparison between mosquitoes at the coolest and warmest temperatures (27°C vs. 32°C), using the cooler temperature as the baseline. To identify the main effect of aging for each immune treatment, irrespective of temperature, we conducted three pairwise DE comparisons: overall aging (1 day vs. 15 days), early aging (1 day vs. 5 days) and late aging (10 days vs. 15 days), using the younger age in each comparison as the baseline.

For each DE comparison, GSEA was performed to identify which biological functions (“Biological Process” GO terms) were enriched, using normalized enrichment scores (NES), where a larger magnitude indicates greater enrichment. A positive NES indicates that genes within the GO term are upregulated, whereas a negative NES indicates that genes within the GO term are downregulated. We next used REVIGO, with *Drosophila* as the closest reference, to summarize the resulting significant GO terms (log_2_ fold change ≥ 1 or ≤ -1; adjusted *p* < 0.05) into common representative terms (64), and parsed them into 12 overarching functional categories: cell adhesion, cell cycle, DNA replication and repair, immunity, metabolism and mitochondrial function, molecule transport, protein processing, transcription, translation and RNA processing, sensory perception, synaptic activity, and miscellaneous.

### Warmer temperature and aging interaction analysis

To identify the interactive effects of warmer temperature and aging, we separated the data by immune treatment and conducted two separate interactive analyses. For each interactive analysis, we separated each temperature-age combination such that there were 12 different conditions (3 temperatures x 4 ages = 12 conditions; e.g. 27°C-1 day, 30°C-5 day, etc.). We conducted all possible pairwise contrasts by DE analysis using DESeq2 (Additional File 4: Table S1) to identify DEGs affected by aging at a specific temperature and DEGs affected by temperature at a specific age. Interaction DEGs were identified as (i) genes that were differentially expressed with aging at one temperature, and this differential expression was different in terms of direction or magnitude at another temperature, or (ii) genes that were differentially expressed with warmer temperature at one age, and this differential expression was different in terms of direction or magnitude at another age (Additional Files 5-7: Files S3-S5).

To infer how the interaction between temperature and age shapes physiology, we performed k-means clustering to analyze expression pattern similarities (65). This analysis grouped the interaction-DEGs into 10 clusters that were each assigned a number as a label. We generated a supervised heatmap of the clustered interaction-DEGs by scaling the mRNA abundance for each gene across all samples to generate a z-score (Additional File 7: File S5). Specifically, Z-scores were generated by subtracting the mean mRNA abundance value across all genes from the individual gene mRNA abundance value, then dividing this difference by the standard deviation of expression. GSEA was then conducted for each cluster to identify which biological functions were enriched (Additional File 8: File S6). Enriched GO terms were then assigned to the 12 functional categories (as described above for main effect GSEAs) to summarize the interactive effects of warmer temperature and aging on biological function. Next, we selected a subset of interaction DEGs that (i) were annotated in the genome with known functions related to immunity, metabolism, or DNA repair, and (ii) demonstrated different patterns of transcriptomic changes due to the interaction between warmer temperature and aging. To visualize these specific examples, we plotted the mRNA abundance of the DEGs.

## Results

### Immune induction, warmer temperature, and aging shape the transcriptome

To determine how immune induction, warmer temperature, aging, and the interaction between temperature and age alter gene expression, we reared mosquitoes at 27°C, 30°C or 32°C, and sequenced the transcriptome of naïve and immune-induced mosquitoes at 1-, 5-, 10-, or 15-days post eclosion (Fig. 1A). To visualize variation in the transcriptome, we performed principal component analysis (PCA) and identified that immune induction, temperature, and age all shape the transcriptome, but immune induction and age explain the majority of the variation among mosquitoes (Fig. 1B). Mosquitoes clustered by immune treatment, regardless of their rearing temperature or age, which indicates that mounting an immune response induces global transcriptomic changes. Importantly, mosquitoes of different ages clustered separately, irrespective of immune treatment or temperature, with 1-day-old mosquitoes being markedly different from any other age. Mosquitoes reared at different temperatures also formed clusters, irrespective of immune treatment or age, although these clusters were less distinct, indicating that temperature has less of an effect on variation in the transcriptome. In summary, immune induction, warmer temperature, and aging shape the mosquito transcriptome.

**Fig 1.**
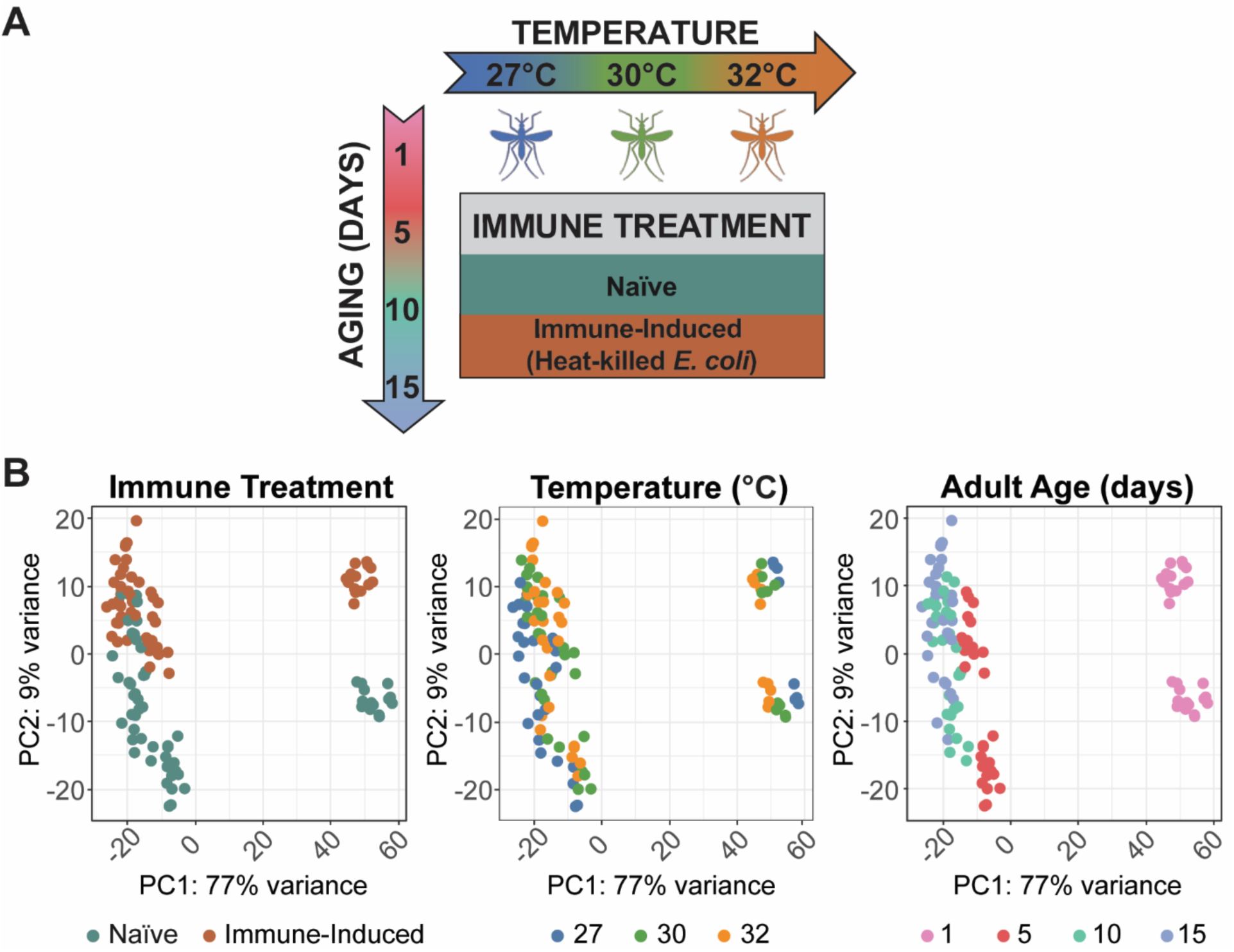
Immune treatment, temperature, and adult age shape the mosquito’s transcriptome. **(A)** Experimental overview to assess the individual and interactive effects of warmer temperature (27°C, 30°C, or 32°C) and aging (1, 5, 10, or 15 days) on the transcriptome of naïve and immune-induced *Anopheles gambiae* mosquitoes. **(B)** Principal component analysis across all samples, color-coded by immune treatment (left panel), temperature (middle panel), or age (right panel).

### Distinct functional categories of the transcriptome are affected by immune induction, warmer temperature, and aging

To identify broad patterns in the transcriptome, we performed iterative weighted gene co-expression network analysis (iterative WGCNA (59)) across all samples to identify genes that have expression patterns that correlate with each trait: immune induction, temperature or age. We identified 22 modules of similarly co-expressed genes (numbered 1-22), which further clustered into 6 larger interconnected metamodules (lettered A through F) (Fig. 2A).

**Fig 2.**
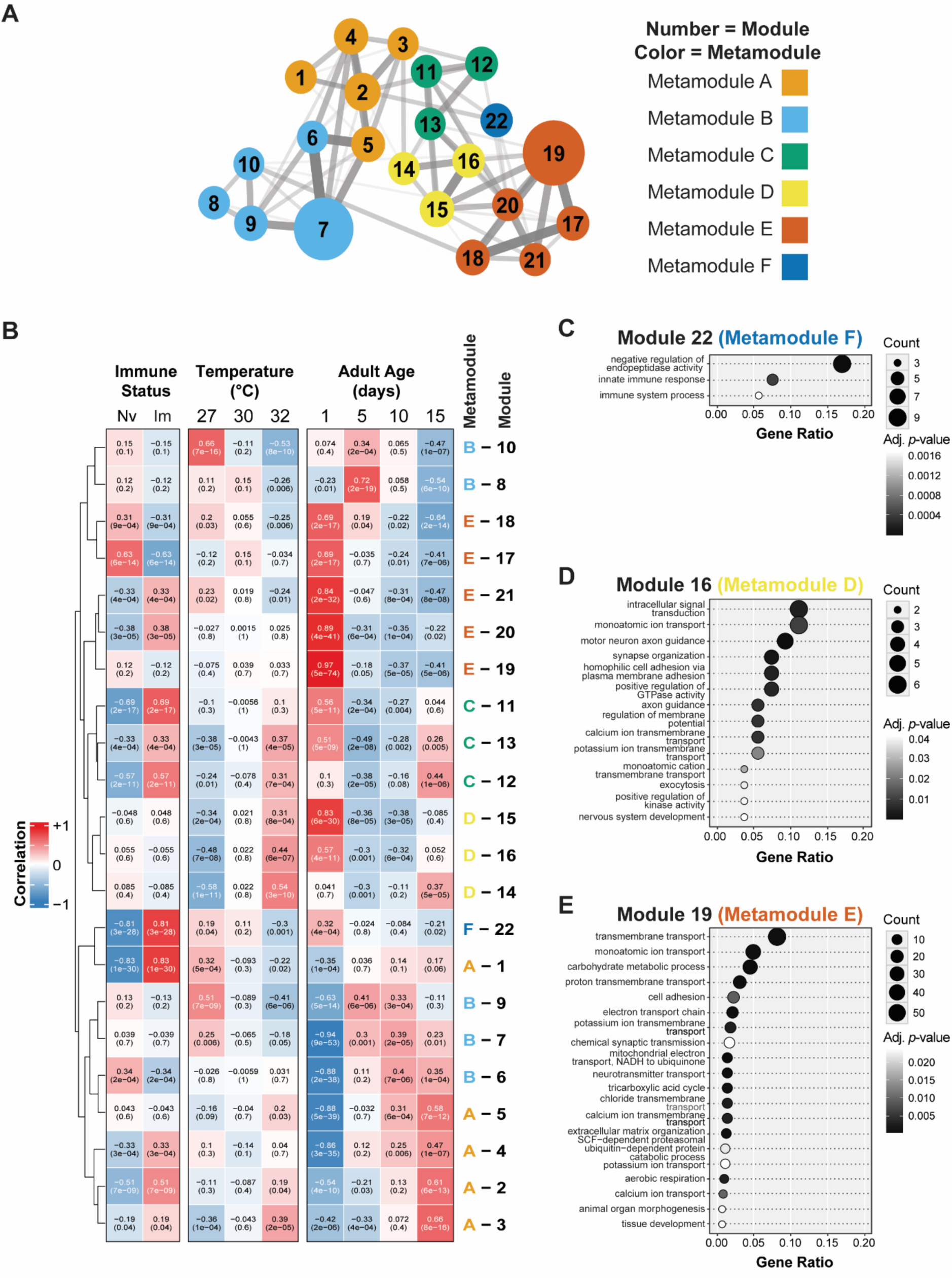
Immune treatment, temperature, and age have unique effects on different functional parts of the mosquito’s transcriptome. **(A)** Diagram illustrating the outcome of iterative weighted gene co-expression network analysis (iWGCNA), which identified 22 modules of interconnected genes (1–22) that further clustered into 6 metamodules (A-F). Lines between modules indicate interconnectivity and the thicker lines indicate a stronger connectivity. **(B)** Heatmap of trait-correlation analysis illustrating the correlation between each module and each trait (e.g. temperature) at each level of the trait (e.g. 27°C, 30°C, or 32°C). Naïve and immune-induced levels of the immune status trait are abbreviated as Nv and Im, respectively. Each box depicts the correlation value on the first line and its *p*-value on the second line. Positive correlations are red and negative correlations are blue, with stronger correlations having more intense colors. Modules were arranged using unsupervised hierarchical clustering based on Euclidian distances, with shorter distances indicating more similar expression patterns. **(C-E)** Over-representation analysis of Module 22 **(C)**, Module 16 **(D)**, and Module 19 **(E)**, where gene ratio (enrichment) is on the horizontal axis and gene ontology (GO) term is on the vertical axis. Dot size indicates the number of genes enriched within each category and dot color indicates the p-value (adjusted *p*<0.05).

We then determined how genes in each module correlate with each level within a trait by performing a module-trait-correlation analysis (Fig. 2B). A strong correlation indicates how, for a trait (e.g., temperature), gene expression is shaped by each level within the trait (e.g., 27°C, 30°C or 32°C); this correlation can either be positive (a general increase in expression of genes in the module) or negative (a general decrease in expression of genes in the module). We plotted the trait-correlations on a heatmap and conducted over-representation analysis (ORA) to identify enriched gene ontology (GO) terms in each module (Fig. 2B-E). Most modules were shaped by one trait more than by another (Fig. 2B). For example, metamodule F strongly correlates with immune induction, but weakly correlates with temperature or age. Metamodule F is the only metamodule that is composed of one module, and that module, module 22, positively correlates with genes involved in immunity and the negative regulation of endopeptidase activity (Fig. 2C). Metamodule D strongly correlates with 32°C and 1-day-old mosquitoes but weakly correlates with immune induction. Within metamodule D, module 16 is enriched for genes involved in cell signaling, molecule transport, and synaptic activity (Fig. 2D). Metamodule E strongly correlates with 1-day-old mosquitoes, moderately correlates with immune induction, and weakly correlates with temperature. Within metamodule E, module 19 is enriched for genes involved in metabolism and mitochondrial function and molecule transport (Fig. 2E). In summary, 22 modules are uniquely shaped by immune induction, warmer temperature, and aging.

### Immune induction upregulates immunity and downregulates metabolism

We next identified the specific genes and biological processes that are shaped by the main effect of immune induction by conducting differential expression (DE) analysis followed by Gene Set Enrichment Analysis (GSEA). Immune induction significantly altered the transcriptome, irrespective of temperature or age. Relative to naïve mosquitoes, immune-induced mosquitoes had 355 differentially expressed genes (DEGs), with 242 genes upregulated and 113 genes downregulated (Table 1). Immune induction upregulated genes involved in immunity and protein processing, but downregulated genes involved in metabolism and mitochondrial function and methylation (DNA replication and repair) (Fig. 3). Additionally, immune induction had a bidirectional effect on genes involved in molecule transport and genes involved in translation and RNA processing. For example, Golgi transport (molecule transport) and tRNA aminoacylation (translation and RNA processing) were upregulated, but ion transport (molecule transport) and rRNA processing (translation and RNA processing) were downregulated. In summary, when a mosquito mounts an immune response, genes involved in immunity and protein processing are upregulated, whereas genes involved in metabolism are downregulated.

**Fig 3.**
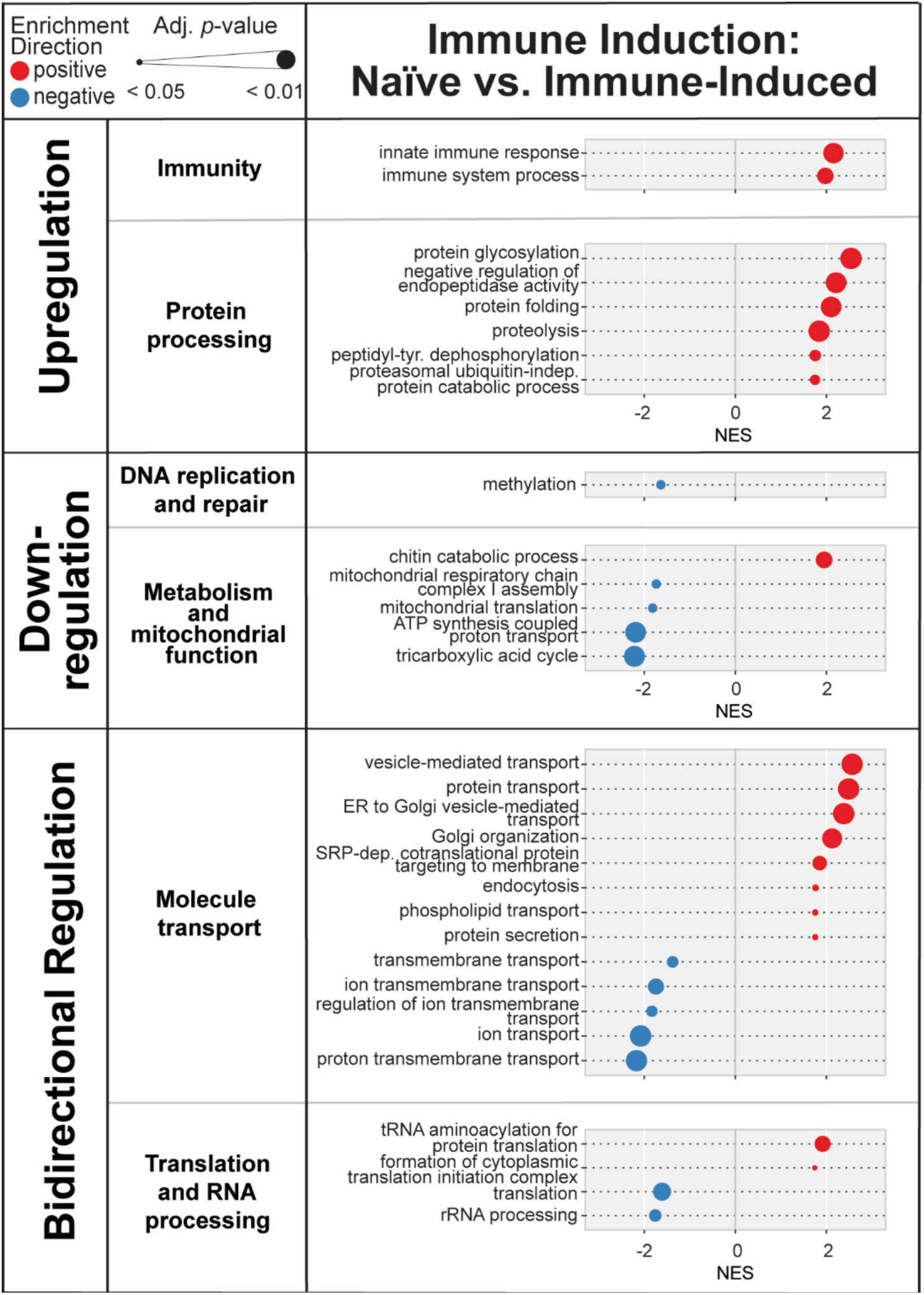
Immune induction upregulates, downregulates or has a bidirectional effect on the expression of genes in distinct functional categories. Diagram illustrates the outcome of Gene Set Enrichment Analysis, where normalized enrichment score (NES) is on the horizontal axis and gene ontology (GO) term is on the vertical axis. Dot sizes indicate the adjusted p-value (*p*<0.05), and dot colors indicate positive enrichment (red) versus negative enrichment (blue).

**Table 1.** Number of genes and biological processes differentially regulated by immune induction.

| Differentially Expressed Genes<br>(Log<sub>2</sub> Fold Change >\|1\|<br>and Adj. *p*<0.05) |  | GSEA |  |
| --- | --- | --- | --- |
|  |  | Number of Functional Categories | Number of Biological Process GO terms |
| Total Number | 355 | 6 | 31 |
| Upregulated | 242 |  |  |
| Downregulated | 113 |  |  |

### Warmer temperature increases metabolism and cell signaling but decreases DNA repair, translation, and protein processing

Because naïve and immune-induced mosquitoes differ across the transcriptome, we next separated these two immune treatments and investigated the main effect of temperature by DE analysis followed by GSEA. For temperature, we compared mosquitoes at 27°C to mosquitoes at 32°C, irrespective of age. Warmer temperature (32°C) resulted in 377 DEGs in naïve mosquitoes and 425 DEGs in immune-induced mosquitoes, with approximately half of the genes being upregulated and half being downregulated (Table 2, Fig. 4).

**Fig 4.**
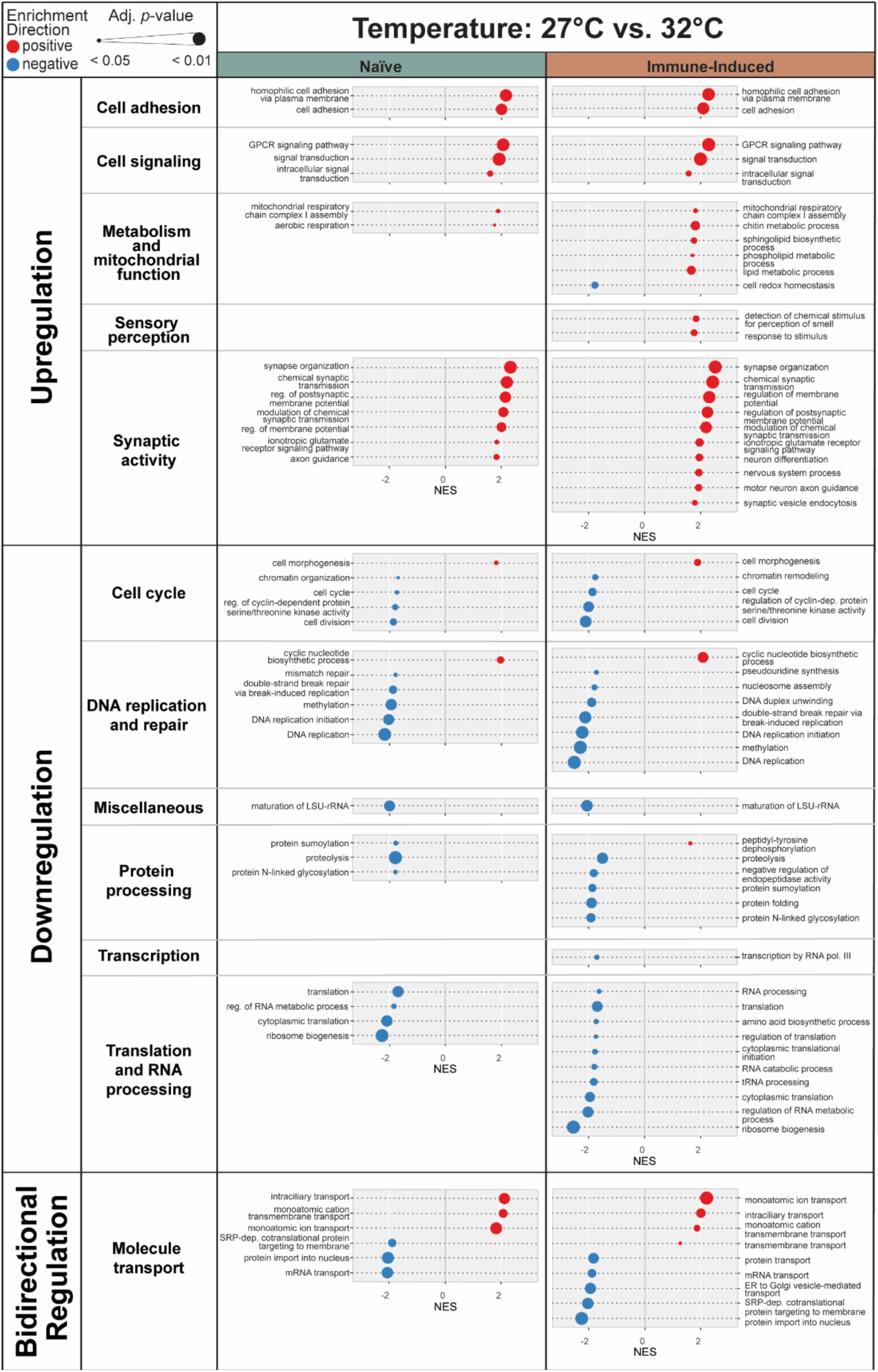
Increasing the temperature from 27°C to 32°C upregulates, downregulates or has a bidirectional effect on the expression of genes in distinct functional categories. Diagram illustrates the outcome of Gene Set Enrichment Analysis, where normalized enrichment score (NES) is on the horizontal axis and gene ontology (GO) term is on the vertical axis. Dot sizes indicate the adjusted p-value (*p*<0.05) and dot colors indicate positive enrichment (red) versus negative enrichment (blue). Left and right gene columns display outcomes in naïve and immune-induced mosquitoes, respectively.

**Table 2.**
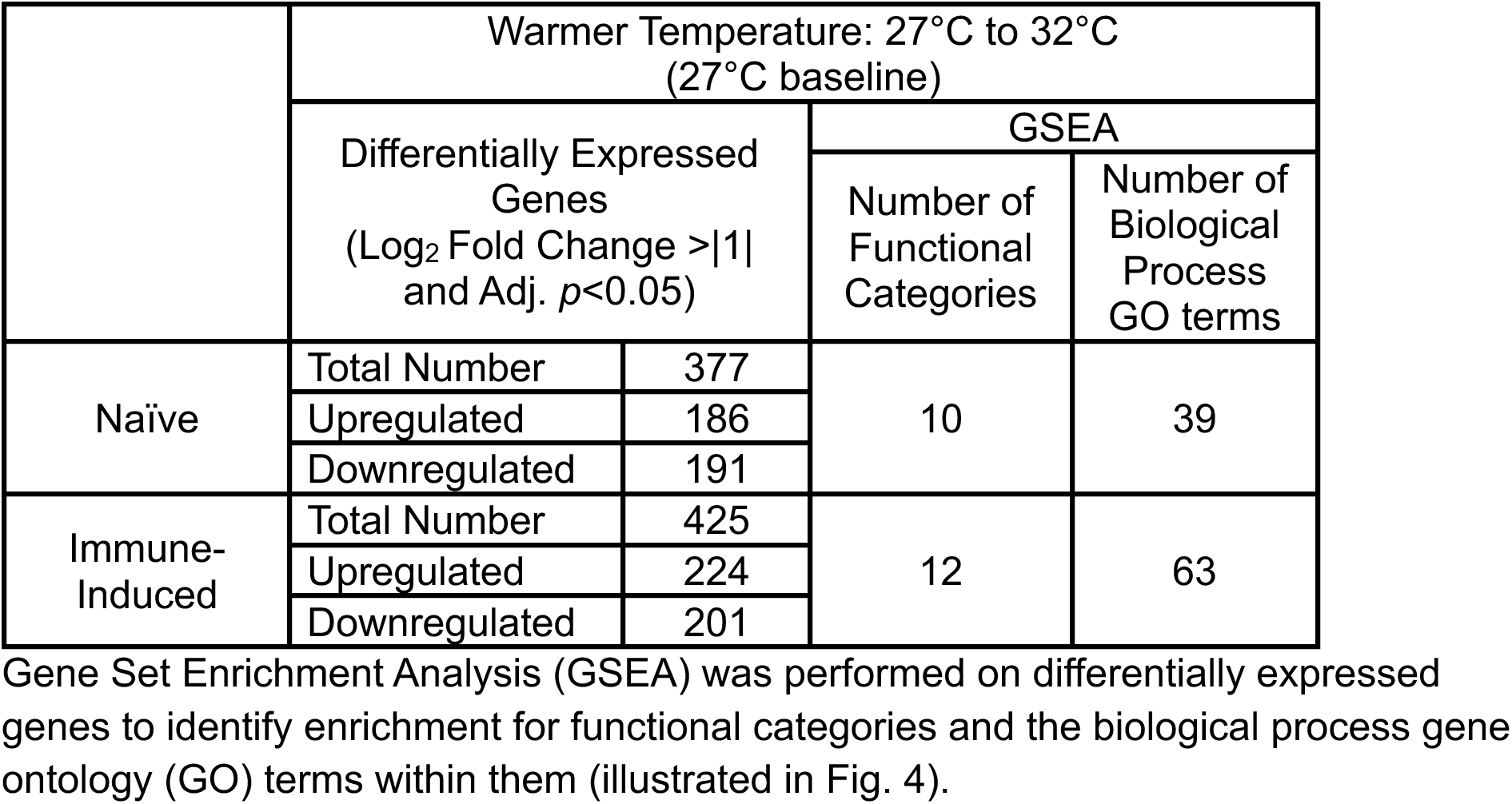
Number of genes differentially regulated when the temperature is increased from 27°C to 32°C.

|  | Warmer Temperature: 27°C to 32°C<br>(27°C baseline) |  |  |  |
| --- | --- | --- | --- | --- |
|  | Differentially Expressed Genes<br>(Log<sub>2</sub> Fold Change >\|1\| and Adj. *p*<0.05) |  | GSEA |  |
|  |  |  | Number of Functional Categories | Number of Biological Process GO terms |
| Naïve | Total Number | 377 | 10 | 39 |
|  | Upregulated | 186 |  |  |
|  | Downregulated | 191 |  |  |
| Immune-Induced | Total Number | 425 | 12 | 63 |
|  | Upregulated | 224 |  |  |
|  | Downregulated | 201 |  |  |

In both naïve and immune-induced mosquitoes, warmer temperature upregulated genes involved in cell adhesion, cell signaling, synaptic activity and metabolism and mitochondrial processes. The warming-based increase in metabolism is likely due to the metabolic rate being faster when the temperature is warmer (72, 73).

In both naïve and immune-induced mosquitoes, warmer temperature downregulated genes involved in the cell cycle, DNA replication and repair, maturation of LSU-rRNA (miscellaneous), protein processing, and translation and RNA processing. This suggests that 32°C disrupts essential processes like cell division and genome maintenance, and this is tied to the warming-based increase in the production of reactive oxygen species, which damages DNA and inhibits its repair (2, 74–84).

In both naïve and immune-induced mosquitoes, warmer temperature had a bidirectional effect on genes involved in molecule transport. For example, warmer temperature upregulated ion transport genes but downregulated genes involved in protein import into the nucleus.

The effect of warmer temperature was mostly similar between naïve and immune-induced mosquitoes, but warmer temperature caused more DEGs and more enriched functional categories (with more GO terms per category) in immune-induced mosquitoes than in naïve mosquitoes (Table 2). Also, unique to immune-induced mosquitoes was that warmer temperature upregulated genes involved in sensory perception and chitin metabolism (metabolism and mitochondrial function) but downregulated genes involved in transcription.

### Aging decreases metabolism but increases DNA replication and repair, transcription, and translation

We next assessed how overall aging (from 1 to 15 days) shapes the transcriptome of naïve and immune-induced mosquitoes, irrespective of temperature. Overall aging resulted in 3085 DEGs in naïve mosquitoes and 3006 DEGs in immune-induced mosquitoes, with about one-third of the genes being upregulated and two-thirds being downregulated (Table 3, Fig. 5). Based on the number of DEGs, overall aging has a much larger effect on gene expression than warmer temperature or immune induction.

**Fig 5.**
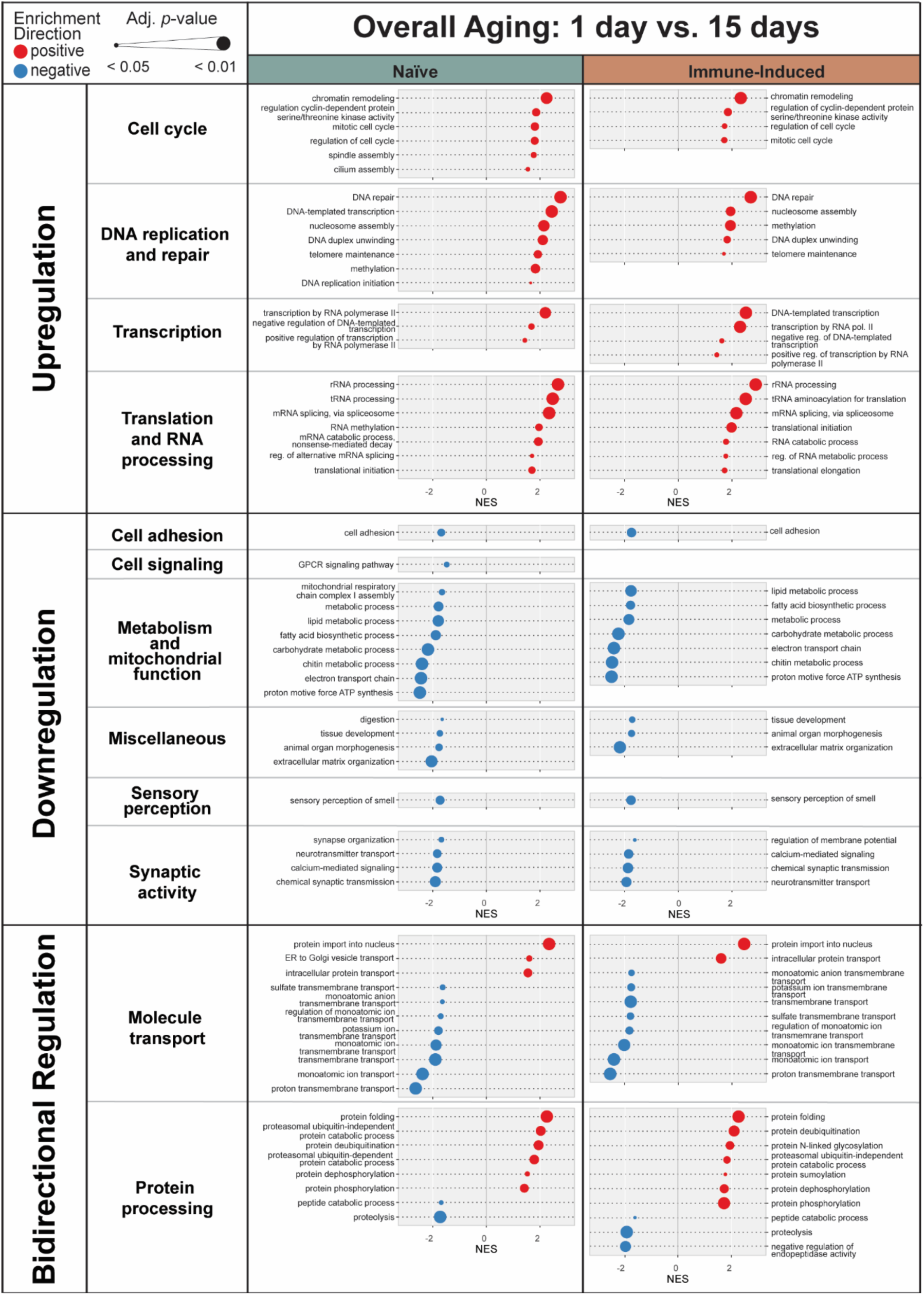
Aging from 1 to 15 days upregulates, downregulates or has a bidirectional effect on the expression of genes in distinct functional categories. Diagram illustrates the outcome of Gene Set Enrichment Analysis, where normalized enrichment score (NES) is on the horizontal axis and gene ontology (GO) term is on the vertical axis. Dot sizes indicate the adjusted p-value (*p*<0.05), and dot colors indicate positive enrichment (red) versus negative enrichment (blue). Left and right gene columns display outcomes in naïve and immune-induced mosquitoes, respectively.

**Table 3.** Number of genes differentially regulated during overall aging (1 to 15 days), early aging (1 to 5 days), and late aging (10 to 15 days) in naïve and immune-induced mosquitoes.

| Age |  | Differentially Expressed Genes<br>(Log<sub>2</sub> Fold Change >\|1\| and Adj. *p*<0.05) |  | GSEA |  |
| --- | --- | --- | --- | --- | --- |
|  |  |  |  | Number of Functional Categories | Number of Biological Process GO terms |
| Overall Aging:<br>1 day to 15 days<br>(1 day is baseline) | Naïve | Total Number | 3085 | 12 | 61 |
|  |  | Upregulated | 1172 |  |  |
|  |  | Downregulated | 1913 |  |  |
|  | Immune-Induced | Total Number | 3006 | 11 | 56 |
|  |  | Upregulated | 1063 |  |  |
|  |  | Downregulated | 1943 |  |  |
| Early Aging:<br>1 day to 5 days<br>(1 day is baseline) | Naïve | Total Number | 2529 | 12 | 68 |
|  |  | Upregulated | 953 |  |  |
|  |  | Downregulated | 2529 |  |  |
|  | Immune-Induced | Total Number | 2476 | 12 | 65 |
|  |  | Upregulated | 880 |  |  |
|  |  | Downregulated | 1596 |  |  |
| Late Aging:<br>10 days to 15 days<br>(10 days is baseline) | Naïve | Total Number | 133 | 10 | 45 |
|  |  | Upregulated | 67 |  |  |
|  |  | Downregulated | 66 |  |  |
|  | Immune-Induced | Total Number | 80 | 10 | 41 |
|  |  | Upregulated | 34 |  |  |
|  |  | Downregulated | 46 |  |  |

In both naïve and immune-induced mosquitoes, overall aging upregulated genes involved in the cell cycle, DNA replication and repair, transcription, and translation and RNA processing. This upregulation of DNA repair is tied to the accumulation of aging-based damage and the aging-based decrease in the efficiency of DNA repair that occurs in dipteran insects (79–84) (85–88).

In both naïve and immune-induced mosquitoes, overall aging downregulated genes involved in cell adhesion, metabolism and mitochondrial function, sensory perception of smell (sensory perception), and synaptic activity. The downregulation of metabolic genes reflects the reduced metabolic rate that is a characteristic of senescence (89–92). Additionally, overall aging downregulated genes involved in tissue development, organ morphogenesis, and extracellular matrix organization (miscellaneous). This is likely because 1-day-old mosquitoes express the genes needed to complete the body reorganization that occurs during hematophagic and gonotrophic capacitation immediately after eclosion (1, 93–96).

In both naïve and immune-induced mosquitoes, overall aging had a bidirectional effect on genes involved in molecule transport and protein processing. Specifically, genes involved in protein import into the nucleus (molecule transport), intracellular protein transport (molecule transport), protein folding (protein processing) and de-ubiquitination (protein processing) were upregulated, whereas genes involved in ion transport (molecule transport) and proteolysis (protein processing) were downregulated.

The effect of overall aging was largely similar between naïve and immune-induced mosquitoes. However, in naïve mosquitoes, but not in immune-induced mosquitoes, aging downregulated cell signaling and digestion (miscellaneous).

### Early and late aging have opposing effects on many biological processes, but both decrease metabolism

Because 1-day-old mosquitoes cluster very differently from all other ages (Fig. 1), we evaluated the effect of aging in naïve and immune-induced mosquitoes, stratified by early aging (from 1 to 5 days) and late aging (from 10 to 15 days). Most changes in gene expression occurred during early aging, regardless of immune treatment or temperature (Table 3, Fig. 6). Specifically, early aging resulted in 2529 DEGs in naïve mosquitoes and 2476 DEGs in immune-induced mosquitoes, with about one-third of the DEGs being upregulated and two-thirds being downregulated. Late aging resulted in 133 DEGs in naïve mosquitoes and 80 DEGs in immune-induced mosquitoes, with approximately half of the DEGs being upregulated and half being downregulated.

**Fig 6.**
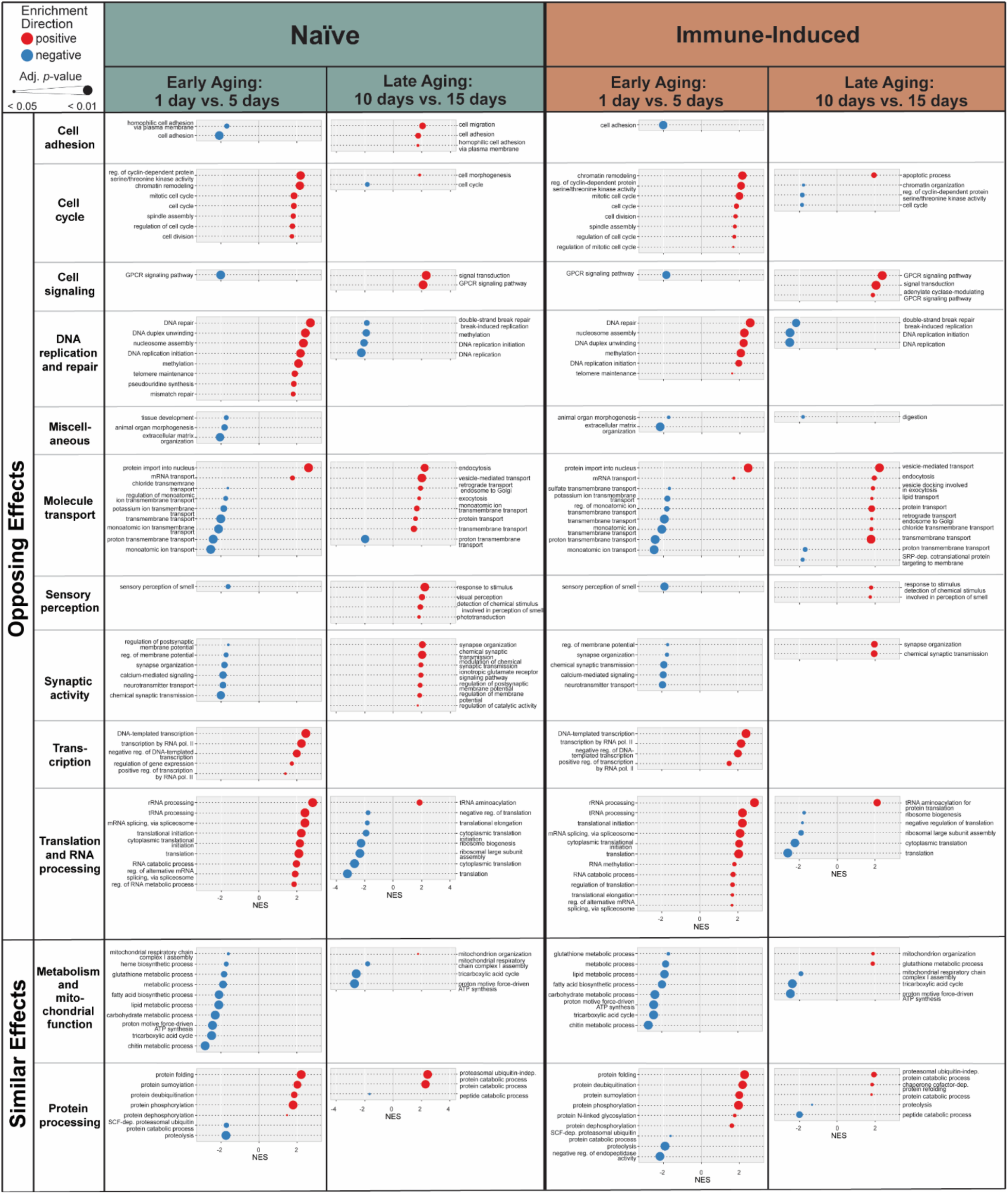
Early aging (1 to 5 days) and late aging (10 to 15 days) have different effects on the regulation of the expression of genes in distinct functional categories. Diagram illustrates the outcome of Gene Set Enrichment Analysis, where normalized enrichment score (NES) is on the horizontal axis and gene ontology (GO) term is on the vertical axis. Dot sizes indicate the adjusted p-value (*p*<0.05) and dot colors indicate positive enrichment (red) versus negative enrichment (blue). Columns display outcomes in naïve (left) and immune-induced (right) mosquitoes, with each column split into early aging and late aging.

In both naïve and immune-induced mosquitoes, early and late aging had opposing effects on many biological processes. Early aging upregulated genes involved in the cell cycle, DNA replication and repair, and translation and RNA processing, but these same processes were downregulated during late aging. In contrast, early aging downregulated genes involved in cell signaling, synaptic activity, and the sensory perception of smell (sensory perception), but these processes were upregulated during late aging. Moreover, early and late aging altered different processes involved in molecule transport.

In naïve and immune-induced mosquitoes, both early and late aging downregulated genes involved in metabolic processes (Fig. 6), which matches the decline in metabolism observed with overall aging (Fig. 5). Additionally, both early and late aging had a bidirectional effect on genes involved in protein processing. Specifically, early aging upregulated protein folding and modification but downregulated proteolysis. Late aging upregulated protein catabolism but downregulated peptide catabolism.

Finally, early aging caused not just more DEGs but also enriched for more biological processes than late aging (Table 3). Unique to early aging was that genes involved in transcription were upregulated, but genes involved in chitin metabolism (metabolism and mitochondrial function) were downregulated. In summary, early and late aging enrich for different biological processes. However, both early and late aging downregulate metabolism, which corresponds to the decline in metabolism that occurs with overall aging.

### The interaction between warmer temperature and aging shapes the transcriptome

Because warmer temperature and aging interact to deteriorate body condition, decrease survival, and weaken reproduction and immunity (1, 9, 12, 42–44), we next queried whether these two traits interact to shape the transcriptome. For this, we conducted differential expression comparisons to identify genes that were (i) differentially expressed with aging at one temperature and this effect changed in magnitude or direction at another temperature, or (ii) differentially expressed with warmer temperature at one age and this effect changed in magnitude or direction at another age. We uncovered that warmer temperature interacts with aging to alter the transcriptome of both naïve and immune-induced mosquitoes (Fig. 7A). This interaction resulted in 5362 DEGs in naïve mosquitoes and 5246 DEGs in immune-induced mosquitoes. Of these DEGs, 794 genes were unique to naïve mosquitoes, 678 genes were unique to immune-induced mosquitoes, and 4568 DEGs were shared.

**Fig 7.**
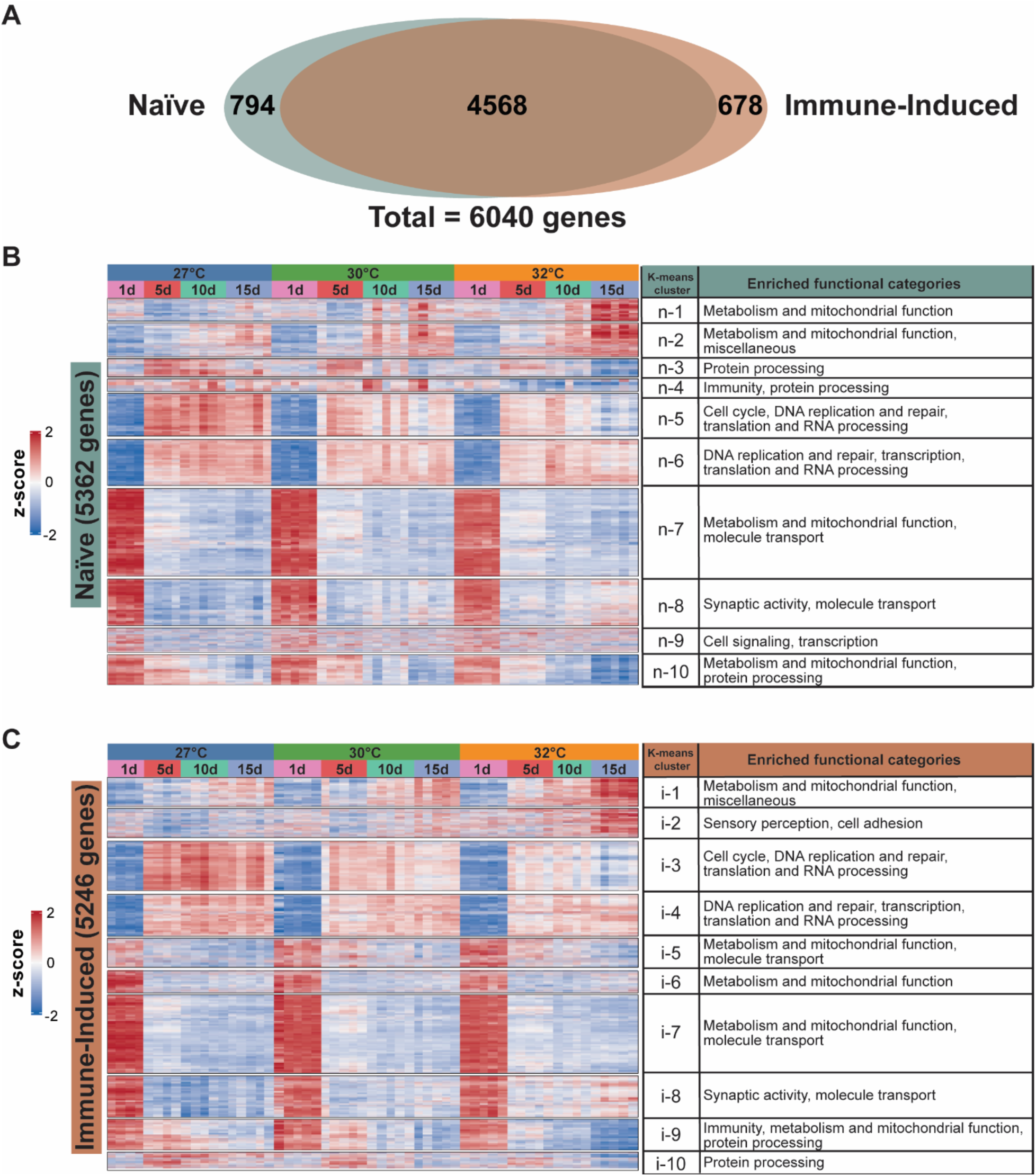
The interaction between warmer temperature and aging shapes biological processes within the mosquito. **(A)** Venn diagram illustrating the number of differentially expressed genes (DEGs) affected by the interaction between temperature and age in naïve and immune-induced mosquitoes. **(B-C)** Heatmaps illustrating mRNA abundance of genes grouped by k-means clustering in naïve **(B)** and immune-induced **(C)** mosquitoes, including the enriched functional categories in each cluster. Interaction-based DEGs were grouped by expression pattern similarity into 10 clusters. Z-scores indicate mRNA abundance scaled across samples, and color indicates abundance: higher is red whereas lower is blue. More intense colors depict larger absolute value of the z-score.

We next characterized how the interaction between temperature and age affects different biological processes by k-means clustering followed by GO enrichment (Fig. 7B and 7C). Notably, in naïve mosquitoes, aging downregulated genes involved in metabolism and mitochondrial function, and when the temperature was warmer, the aging-dependent downregulation occurred earlier in life (k-means cluster n-10) (Fig. 7B). Also, aging upregulated genes involved in the cell cycle, DNA replication and repair, and translation and RNA processing, but this aging-dependent upregulation was smaller when the temperature was warmer (k-means cluster n-5) (Fig. 7B). In immune-induced mosquitoes, aging downregulated genes involved in immunity, metabolism and mitochondrial function, and protein processing, and when the temperature was warmer, the aging-dependent downregulation occurred earlier in life (k-means cluster i-9) (Fig. 7C). In summary, warmer temperature and aging interact to shape the transcriptome.

### Warmer temperature amplifies the aging-dependent decrease in immunity

Many genes are affected by the interaction between warmer temperature and aging (Additional Files 5-7: Files S3-S5). Here, we expand on a few interaction DEGs that are involved in the immune response to infection (Fig. 8, Additional File 9: Table S2).

**Fig 8.**
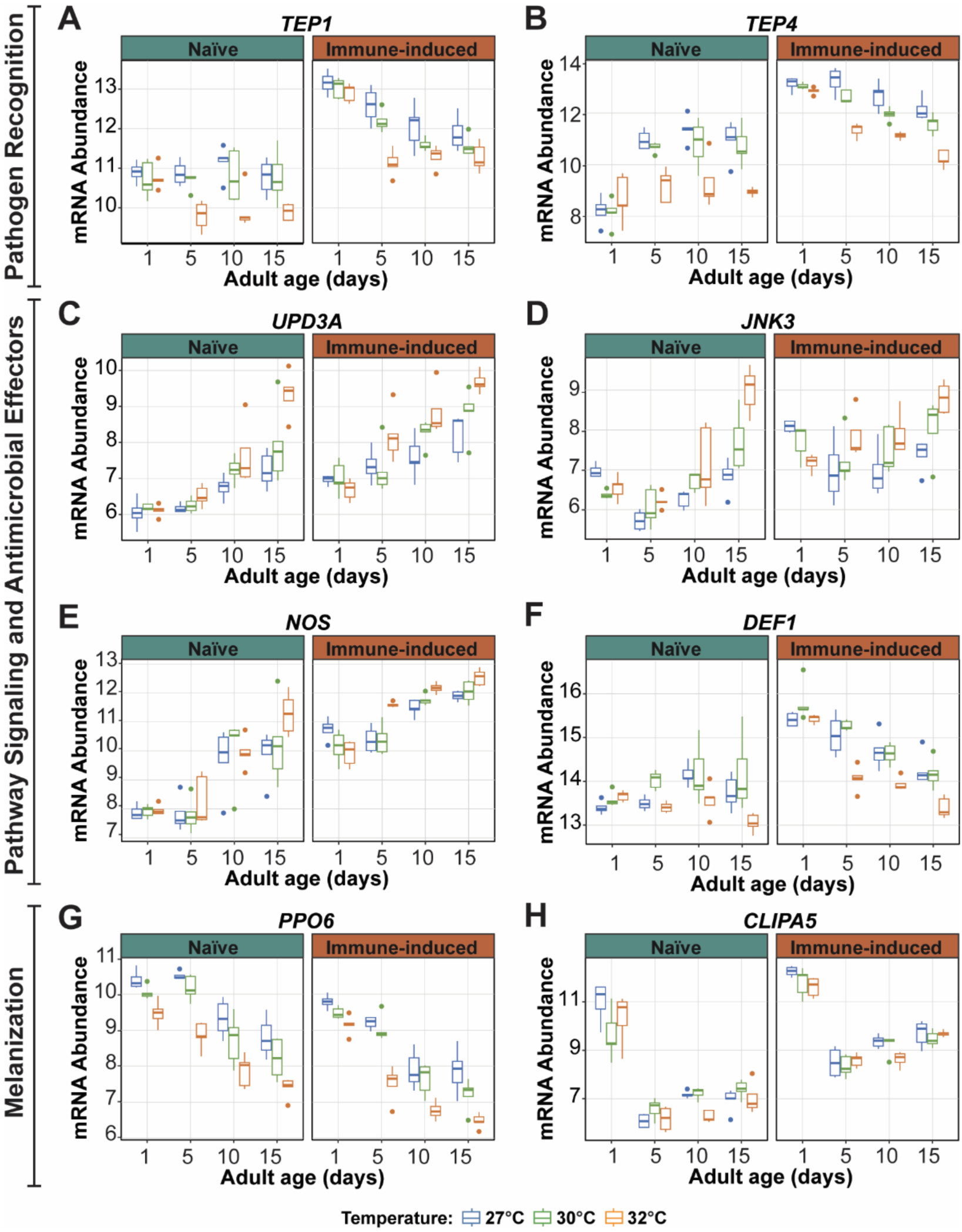
Warmer temperature modifies the aging-dependent changes in immune gene expression. **(A-H)** Variance stabilized transformation (VST) of mRNA abundance illustrating the expression of immune genes in naïve and immune-induced mosquitoes. Box plots display the minimum and maximum, inner quartile range (box), median (horizontal line in box), and outliers (circles). Box plot color depicts rearing temperature. The same information is presented in tabular format in Additional File 9: Table S3.

Complement-like thioester-containing proteins (TEPs) are pattern recognition receptors that positively regulate phagocytosis, melanization, and the infection-induced aggregation of hemocytes on the heart (97–103). In both naïve and immune-induced mosquitoes, *TEP1* and *TEP4* expression in 1-day-old mosquitoes was similar for all temperatures. In naïve mosquitoes, *TEP1* expression remained constant with aging except at 32°C, where it markedly decreased, but *TEP4* expression increased with aging except at 32°C, where expression remained constant. In immune-induced mosquitoes, *TEP1* and *TEP4* expression decreased with aging, and this decrease was amplified at 32°C (Fig. 8A-B).

The Janus kinase-signal transducer and activator of transcription (JAK-STAT) and the c-Jun N-terminal kinase (JNK) pathways are important for the antibacterial, antiplasmodial, and antiviral responses (104–110). In both naïve and immune-induced mosquitoes, warmer temperature and aging increased the expression of the JAK-STAT ligand, Unpaired (*UPD3A*), and the JNK transcription factor, c-Jun N-terminal kinase (*JNK3*) (Fig. 8C and 8D). Importantly, the effects of age depended on the temperature: the aging-dependent increase in expression was much greater when the temperature was warmer.

Nitric oxide is an antimicrobial and signaling molecule that is produced by nitric oxide synthase (NOS). In naïve mosquitoes, the general pattern was that *NOS* expression increased with aging in a manner that was the same for all temperatures, except at 15-days-old, where 32°C caused a disproportionate increase in *NOS* mRNA abundance (Fig. 8E). In immune-induced mosquitoes, *NOS* expression in 1-day-old mosquitoes was the same for all temperatures, but beyond this age, 32°C accelerated the aging-dependent increase in *NOS* expression. Another lytic factor is the antimicrobial peptide, Defensin 1 (DEF1). In naïve mosquitoes, *DEF1* expression marginally increased with aging at 27°C and 30°C but decreased with aging at 32°C (Fig. 8F). In immune-induced mosquitoes, warmer temperature and aging decreased *DEF1* expression, and the aging-dependent decrease occurred faster at 32°C.

Melanization is an effector response against bacteria, parasites, and fungi (111–114). The pro-phenoloxidase, PPO6, is a positive driver of melanization, and the CLIP-domain serine protease, CLIPA5, is a negative regulator of this process. In both naïve and immune-induced mosquitoes, warmer temperature and aging decreased *PPO6* expression, and the aging-dependent decrease occurred faster when the temperature was warmer (Fig. 8G). In both naïve and immune-induced mosquitoes, aging from 1 to 5 days sharply decreased *CLIPA5* expression, whereas aging from 5 to 15 days slightly increased expression, irrespective of temperature (Fig. 8H). In both naïve and immune-induced mosquitoes, warmer temperature did not affect the expression of *CLIPA5* at most ages. However, at 10-days-old, the expression of *CLIPA5* was lowest at 32°C. These gene expression changes explain our phenotypic observation that the aging-based reduction of melanization is faster when the temperature is warmer (12).

### Warmer temperature dampens the aging-dependent decrease in metabolism and the aging-dependent increase in DNA repair

Many metabolic and DNA repair genes are also affected by the interaction between temperature and age (Additional Files 4-6: Files S3-S5). Here, we expand on a few interaction DEGs that are involved in metabolism and DNA repair (Fig. 9, Additional File 10: Table S3).

**Fig 9.**
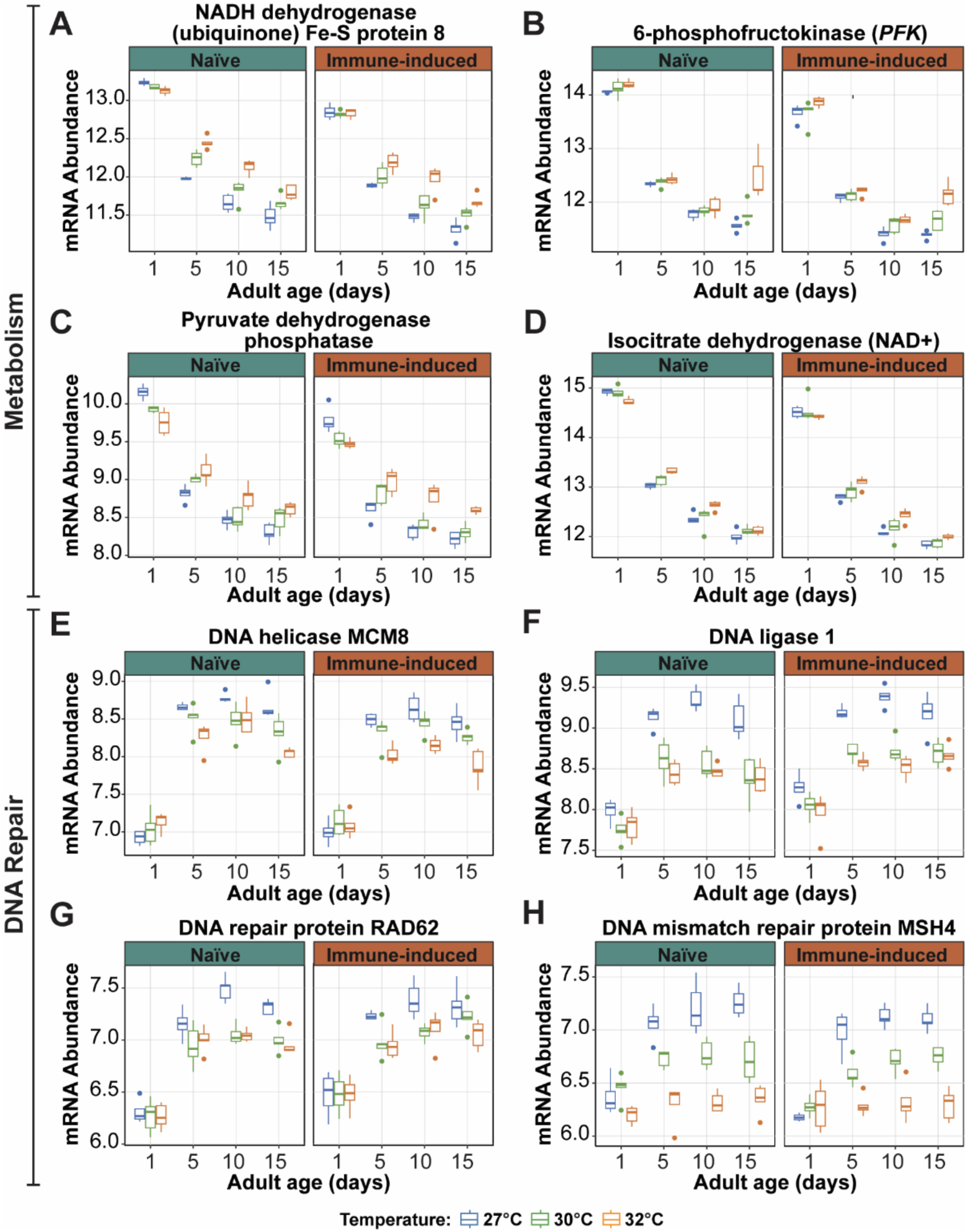
Warmer temperature modifies the aging-dependent changes in the expression of genes involved in metabolism and DNA repair. **(A-H)** Variance stabilized transformation (VST) of mRNA abundance illustrating the expression of genes involved in metabolism and DNA repair in naïve and immune-induced mosquitoes. Box plots display the minimum and maximum, inner quartile range (box), median (horizontal line in box), and outliers (circles). Box plot color depicts rearing temperature. The same information is presented in tabular format in S2 Table.

The electron transport chain, glycolysis, and the citric acid cycle generate cellular energy in the form of ATP. NADH dehydrogenase is part of complex I in the electron transport chain and transfers electrons from NADH to coenzyme Q (115, 116). Aging decreased the expression of NADH dehydrogenase, but warmer temperature increased expression (Fig. 9A). However, in 1-day-old mosquitoes, the warming-based increase did not occur, and instead, NADH dehydrogenase expression was the same at all temperatures.

ATP-dependent 6-phosphofructokinase (PFK) regulates glycolysis. Aging, especially between 1 and 5 days, decreased *PFK* expression, and warmer temperature caused a small increase (Fig. 9B). However, in 15-day-old mosquitoes, the warming-based increase in *PFK* expression was amplified at 32°C.

Pyruvate dehydrogenase phosphatase regulates the complex that converts pyruvate into acetyl-CoA, which then enters the citric acid cycle. Later, isocitrate dehydrogenase (NAD+) catalyzes the rate-limiting step to produce NADH (117). Aging decreased the expression of both genes whereas warmer temperature generally increased expression (Fig. 9C and 9D). The exception to this pattern was 1-day-old mosquitoes, where 32°C decreased expression.

The ability to repair DNA is critical for survival (87). DNA repair enzymes and proteins maintain chromosomal stability and facilitate mismatch repair, homologous recombination, and alternative-non-homologous end joining (118–120). Aging from 1 to 5 days increased the expression of DNA helicase MCM8 and DNA ligase 1, but further aging stabilized the expression. Expression of *MCM8* and DNA ligase 1 in 1-day-old mosquitoes was similar for all temperatures, but with aging, expression of these genes was lower when the temperature was warmer (Fig. 9E and 9F). A similar phenomenon was observed for the DNA repair genes *RAD62* and *MSH4* (Fig. 9G and 9H).

In summary, expression of metabolic genes decreases with aging, but warmer temperature dampens the aging-based decrease. Conversely, expression of DNA repair genes increases with aging, but warmer temperature dampens the aging-based increase.

## Discussion

Using an RNA-sequencing and network analysis approach, we demonstrate that the transcriptome of mosquitoes is individually and interactively shaped by environmental temperature and aging: warmer temperature modifies the aging-dependent changes in the transcriptome of both naïve and immune-induced mosquitoes (Fig. 10).

**Fig 10.**
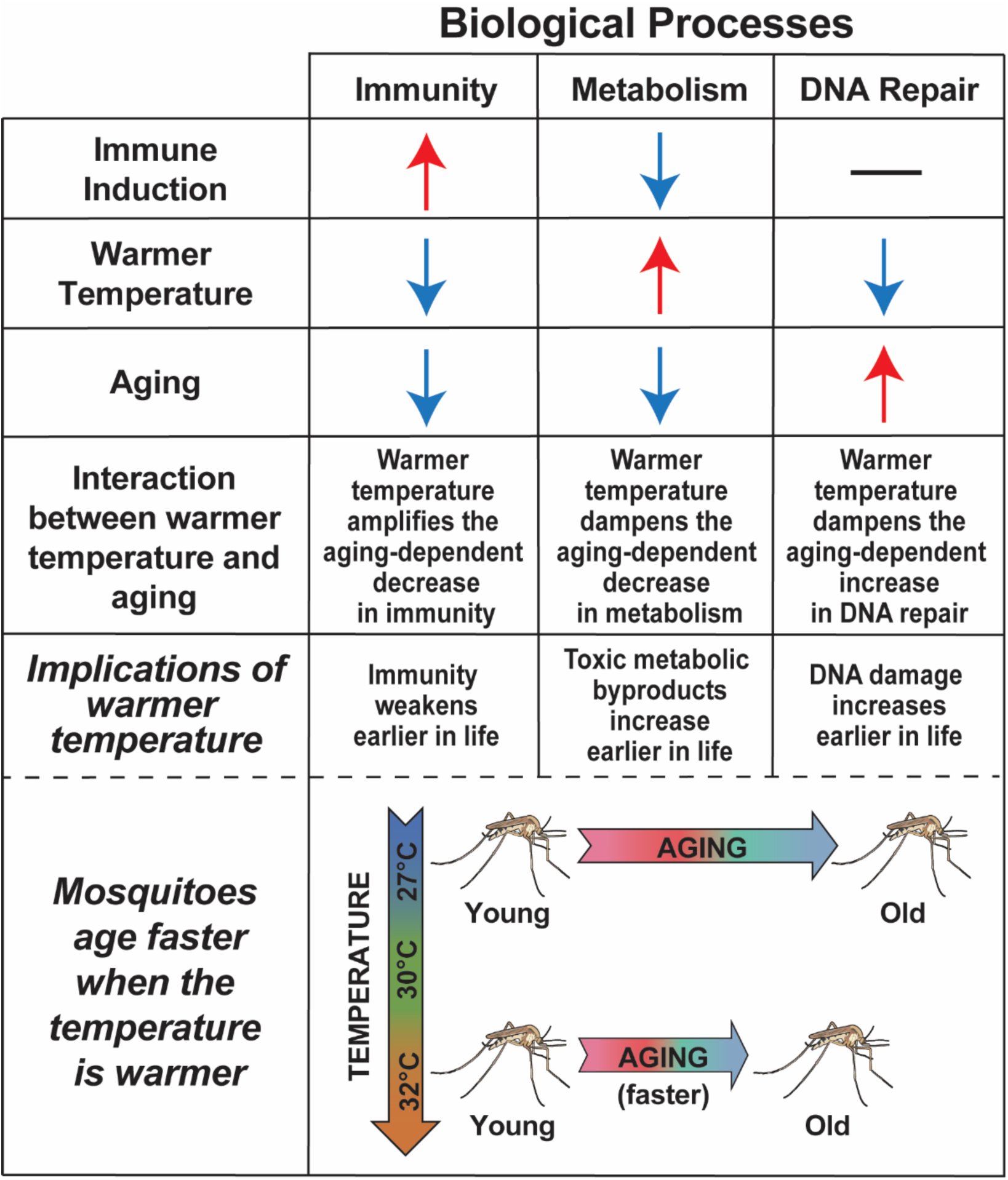
Warmer temperature modifies the aging-dependent changes in the mosquito transcriptome. Red arrows indicate upregulation, blue arrows indicate downregulation, and black dash indicates no change.

Immune induction caused broad changes to the transcriptome of mosquitoes. The upregulation of immunity genes after immune induction was expected. Less anticipated was the downregulation of genes involved in metabolism, which suggests that there is a trade-off between immunity and metabolism. Indeed, mounting an immune response is energetically costly (66–68). Life history theory explains that organisms allocate their finite energetic resources toward processes that increase reproductive output and survival (69, 70). In fruit flies, the activation of encapsulation (an immune response) is linked to a lower metabolic rate (71). This matches our discovery that immune induction upregulates genes involved in immunity while downregulating genes involved in metabolism.

A pronounced discovery was that, regardless of immune status or temperature, aging from 1 to 15 days—and mostly aging from 1 to 5 days—dramatically changes the transcriptome. Aging is known to downregulate the expression of genes involved in metabolism, autophagy, and the maintenance of cuticular structure (35–38). Moreover, older mosquitoes have a reduced metabolic rate and a weakened immune system, which are hallmarks of senescence in Insecta (83, 90, 121). Here, we observed that aging downregulates expression of genes involved in metabolism and immunity but upregulates genes involved in DNA repair. For example, aging decreases expression of NADH dehydrogenase (metabolism) and *PPO6* (immunity) but increases expression of *MCM8* (DNA repair). An aging-based decrease in NADH dehydrogenase mRNA abundance correlates with the aging-based decrease in NADH dehydrogenase protein abundance, and the aging-based decrease in *PPO6* mRNA abundance correlates with the aging-based weakening of the melanization immune response (12, 39, 92). Moreover, the aging-dependent increase in mRNA abundance of *MCM8* and other DNA repair genes is likely due to the aging-dependent accumulation of DNA damage, which requires repair to maintain fidelity and chromosomal integrity (85–88). Importantly, the aging-dependent changes we observed occurred at all temperatures investigated, indicating that aging greatly alters physiology regardless of environmental temperature.

Notably, early and late aging had opposing effects on many genes and biological processes. Most aging-based changes in the transcriptome occurred during early aging, making the transcriptomic profile of 1-day-olds very different from all other ages. For example, the expression of *PFK* (metabolism) was much higher in 1-day-olds than in any other age. This and other changes are likely explained by 1-day-olds having a higher metabolic rate while carrying out tissue remodeling and developmental processes over the first 3 days of adulthood that result in sexual maturity (1, 93–96, 122, 123). Moreover, this marked difference with early aging matches findings in *Aedes aegypti* and *Aedes albopictus*, where 1-day-old mosquitoes have a much higher expression of genes involved in calcium binding and maintenance of cuticular structure, but a much lower expression of genes involved in cell division compared to older mosquitoes (124).

Environmental temperature also modifies the transcriptome. Warmer temperature quickens metabolism, weakens immunity, and lowers survival (2–5, 7, 9, 10, 12, 13, 16–18). Here, we observed that warmer temperature upregulates processes like metabolism and cell signaling but downregulates processes like DNA repair and protein processing. For example, warmer temperature upregulates the expression of NADH dehydrogenase (metabolism) but downregulates the expression of *PPO6* (immunity) and *MCM8* (DNA repair). Similar changes in gene expression with warmer temperature have also been observed in *Aedes aegypti* and *Anopheles stephensi*, with warmer temperature having a global effect on the mosquito transcriptome (16–18). Importantly, the temperature-dependent transcriptomic changes we observed occurred even as the mosquito aged, demonstrating that warmer temperature alters biological processes regardless of age.

We discovered that warmer environmental temperature modifies the effects of aging on the transcriptome. Warmer temperature accelerates senescence in mosquitoes, deteriorating body condition, decreasing reproduction, weakening immunity, and reducing survival (1, 9, 12, 42–46). In *D. melanogaster*, *C. elegans*, and even humans, warmer environmental temperature reduces longevity, likely accelerating the onset of senescence (9, 15, 125–128). Here, we observed that warmer temperature and aging interact to shape the expression of over 6000 genes, broadly altering immunity, metabolism and mitochondrial function, the cell cycle, protein processing, sensory perception, DNA replication and repair, and other processes. For example, warmer temperature amplifies the aging-dependent decrease in the expression of *PPO6* (immunity), weakening the melanization immune response. Additionally, warmer temperature dampens the aging-dependent decrease in the expression of metabolic genes that generate ATP through the electron transport chain, glycolysis, and the citric acid cycle. This is likely because mosquitoes have an overall faster metabolic rate when the temperature is warmer (14, 15, 129). Finally, warmer temperature dampens the aging-based increase in the expression of DNA repair genes, and this correlates with warmer temperature inhibiting DNA repair (2, 74–78, 130).

The goal of our study was to capture transcriptional effects associated with the entire life history of the mosquito. We reared mosquitoes from egg to adulthood at the temperature of experimentation—27°C, 30°C or 32°C—and did so because the experiences of the immature stages can have effects that carry over into the adult stage (131–136). In general, adults that eclose from larvae reared at warmer temperature are smaller than adults that eclose from larvae reared at cooler temperature (4, 11, 137, 138). However, these studies on adult size have predominantly examined temperature ranges that are wider than the ones assessed here. In a prior study, we reared mosquitoes at 27°C, 30°C and 32°C, and found only marginal changes in the size of the resulting adults; wing length was unchanged, and tibia and abdomen length was only 4% larger in mosquitoes reared at 27°C than in mosquitoes reared at 32°C (1). So, we infer that the transcriptional changes observed in the present study are not impacted by adult size. However, gene expression may be impacted by other aspects of life history. For example, the protein content of sugar fed mosquitoes decreases at warmer temperature and with aging, and the aging-associated decrease in protein content accelerates when the temperature is warmer (1). Overall, this study provides an accurate reading of how temperature, aging and their interaction influence the transcriptome in a natural environment, where the temperature in which the immatures develop equates to the temperature in which the adults inhabit.

## Conclusions

In summary, warmer temperature and aging individually alter the transcriptome, and importantly, warmer temperature modifies the aging-dependent changes. In both naïve and immune-induced mosquitoes, warmer temperature amplifies the aging-dependent decrease in immunity but dampens both the aging-dependent decrease in metabolism and the aging-dependent increase in DNA repair. Altogether, warmer temperature accelerates senescence, dramatically shaping the transcriptome of mosquitoes in ways that reflect changes in their ability to respond to infection and survive in our warming world.

## Declarations

### Ethics approval and consent to participate

Not applicable.

### Consent for publication

Not applicable.

### Availability of data and materials

The RNAseq data supporting the conclusions of this article are accessible through GEO Series accession number GSE284722. Additional analyses supporting the conclusions of this article are included in the main text and its additional files.

### Competing interests

The authors declare no competing interests.

### Funding

This work was funded by National Science Foundation (NSF) Grant 1936843 to JFH and NSF Graduate Research Fellowship to LEM.

### Authors’ contributions (CRediT)

Conceptualization: LEM, JSB, JFH. Data Curation: LEM, JSB, J-PC, SS, TYE-L, JFH. Formal analysis: LEM, JSB, J-PC, SS, JFH. Funding acquisition: LEM, JFH. Investigation: LEM, JSB, TYE-L. Methodology: LEM, JSB, J-PC, SS, TYE-L, JFH. Supervision: LEM, JSB, JFH. Visualization: LEM, JSB, J-PC, SS, JFH. Writing – original draft: LEM, JSB, JFH. Writing – review and editing: LEM, JSB, J-PC, SS, TYE-L, JFH.

## Acknowledgements

We thank Drs. Ann Tate, Courtney Murdock, and Leah Sigle for their helpful advice and discussion about RNAseq study design. We thank Cole Meier, Saksham Saksena, Seokin Yang, and Puspa Shah for valuable discussions in the preparation of the manuscript. We thank Megan Grant and Shabbir Ahmed for their valuable commentary on the manuscript draft. The mosquito images depicted in Fig 10 were obtained from the NIH BioArt Source (https://bioart.niaid.nih.gov) and modified.

## Notes

### Competing Interest Statement

The authors have declared no competing interest.

