## Additional File 1 for "Warmer temperature accelerates senescence by modifying the aging-dependent changes in the mosquito transcriptome, altering immunity, metabolism, and DNA repair"

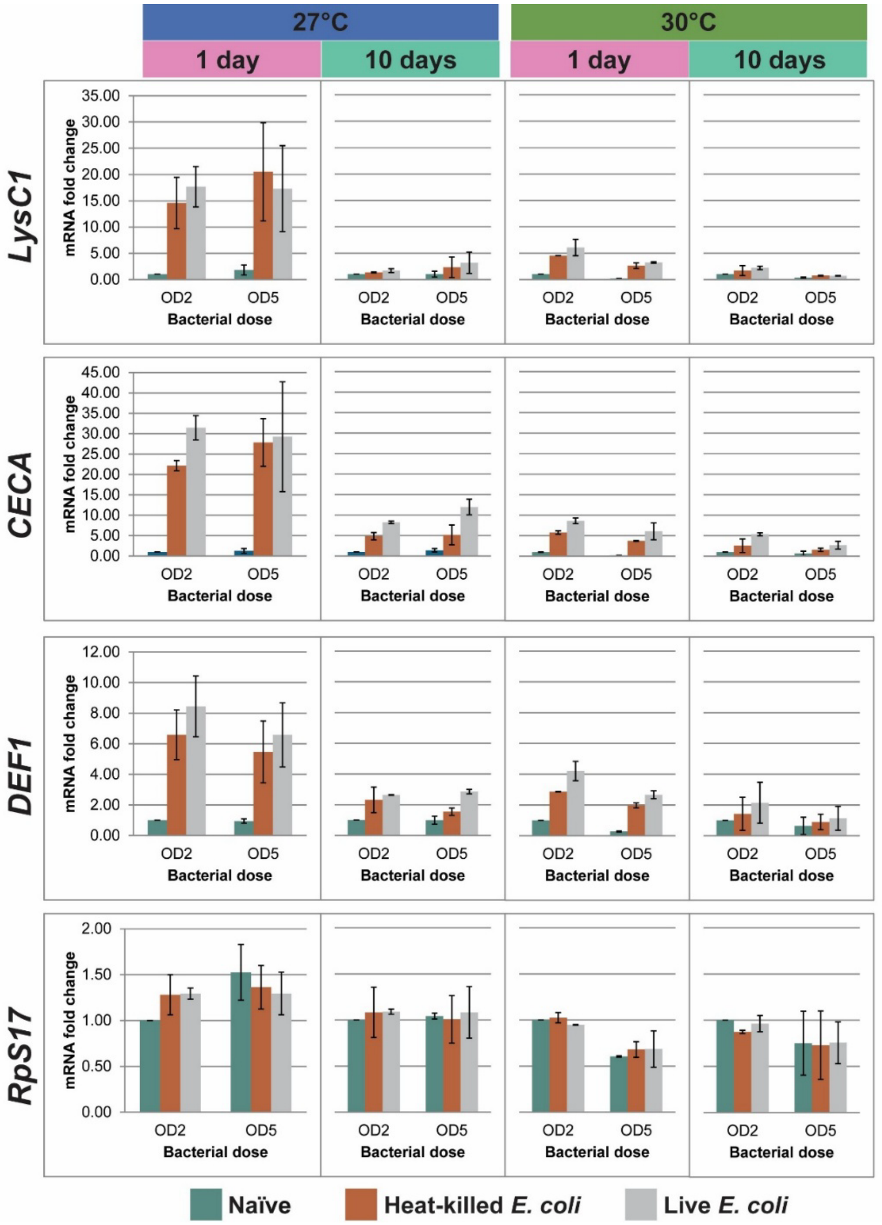


**Fig. S1. Preliminary experiments designed to select the type of immune treatment and dose show that immune gene expression at 6 h post immune treatment does not differ between live or heat-killed *E. coli* injection, nor between bacterial doses.** The relative abundance of mRNA of three immune genes (*LysC1*, *CECA*, and *DEF1*) and one house keeping gene (*RPS17*) was measured by qRT-PCR using the $2^{-\Delta\Delta CT}$ method and *RpS7* as the reference. Heat-killed and live bacterial doses included a low dose ($OD_{600}$ = 2) and a high dose ($OD_{600}$ = 5). Primers used and methods for cDNA synthesis and qRT-PCR are as we have described (https://doi.org/10.1186/s13071-017-2302-6). Data show two independent biological trials. Column heights represent mean fold change with error bars indicating the standard deviation. Rearing temperature and mosquito age are depicted at the top of the plots.
