## Additional File 4 for "Warmer temperature accelerates senescence by modifying the aging-dependent changes in the mosquito transcriptome, altering immunity, metabolism, and DNA repair"

**Table S1. List of pairwise contrasts used to determine interactive effects of warmer temperature and aging for naïve and immune-induced mosquitoes.**

| Naïve Contrasts | | | Immune-Induced Contrasts | | |
|---|---|---|---|---|---|
| Temperature effect in each age | | | Temperature effect in each age | | |
| Contrast # | Comparison Group | Baseline Group | Contrast # | Comparison Group | Baseline Group |
| 1 | 5d_30C | 5d_27C | 31 | 5d_30C | 5d_27C |
| 2 | 5d_32C | 5d_27C | 32 | 5d_32C | 5d_27C |
| 3 | 5d_32C | 5d_30C | 33 | 5d_32C | 5d_30C |
| 4 | 1d_30C | 1d_27C | 34 | 1d_30C | 1d_27C |
| 5 | 1d_32C | 1d_27C | 35 | 1d_32C | 1d_27C |
| 6 | 1d_32C | 1d_30C | 36 | 1d_32C | 1d_30C |
| 7 | 10d_30C | 10d_27C | 37 | 10d_30C | 10d_27C |
| 8 | 10d_32C | 10d_27C | 38 | 10d_32C | 10d_27C |
| 9 | 10d_32C | 10d_30C | 39 | 10d_32C | 10d_30C |
| 10 | 15d_30C | 15d_27C | 40 | 15d_30C | 15d_27C |
| 11 | 15d_32C | 15d_27C | 41 | 15d_32C | 15d_27C |
| 12 | 15d_32C | 15d_30C | 42 | 15d_32C | 15d_30C |
| Age effect in each temperature | | | Age effect in each temperature | | |
| Contrast # | Comparison Group | Baseline Group | Contrast # | Comparison Group | Baseline Group |
| 13 | 5d_27C | 1d_27C | 43 | 5d_27C | 1d_27C |
| 14 | 10d_27C | 1d_27C | 44 | 10d_27C | 1d_27C |
| 15 | 15d_27C | 1d_27C | 45 | 15d_27C | 1d_27C |
| 16 | 10d_27C | 5d_27C | 46 | 10d_27C | 5d_27C |
| 17 | 15d_27C | 5d_27C | 47 | 15d_27C | 5d_27C |
| 18 | 15d_27C | 10d_27C | 48 | 15d_27C | 10d_27C |
| 19 | 5d_30C | 1d_30C | 49 | 5d_30C | 1d_30C |
| 20 | 10d_30C | 1d_30C | 50 | 10d_30C | 1d_30C |
| 21 | 15d_30C | 1d_30C | 51 | 15d_30C | 1d_30C |
| 22 | 10d_30C | 5d_30C | 52 | 10d_30C | 5d_30C |
| 23 | 15d_30C | 5d_30C | 53 | 15d_30C | 5d_30C |
| 24 | 15d_30C | 10d_30C | 54 | 15d_30C | 10d_30C |
| 25 | 5d_32C | 1d_32C | 55 | 5d_32C | 1d_32C |
| 26 | 10d_32C | 1d_32C | 56 | 10d_32C | 1d_32C |
| 27 | 15d_32C | 1d_32C | 57 | 15d_32C | 1d_32C |
| 28 | 10d_32C | 5d_32C | 58 | 10d_32C | 5d_32C |
| 29 | 15d_32C | 5d_32C | 59 | 15d_32C | 5d_32C |
| 30 | 15d_32C | 10d_32C | 60 | 15d_32C | 10d_32C |
