## Additional File 9 for "Warmer temperature accelerates senescence by modifying the aging-dependent changes in the mosquito transcriptome, altering immunity, metabolism, and DNA repair"

**Additional File 9: Table S2**

**Table S2. Warmer temperature and aging interactively shape the expression of genes involved in immunity.** Direction of arrow indicates upregulation, downregulation, or no change.

| | | | | NAÏVE | | | IMMUNE-INDUCED | | |
|---|---|---|---|---|---|---|---|---|---|
| | **Gene (AGAP ID)** | **Interaction K-means Cluster** Naïve | HKEC | Warmer Temp | Aging | Temperature-Age Interaction | Warmer Temp | Aging | Temperature-Age Interaction |
| **PATHOGEN RECOGNITION** | ***TEP1*** (AGAP010815) | n-4 | i-9 | ↓ | – | 32°C reduces expression beyond 1 day of age. | ↓ | ↓ | 32°C reduces expression beyond 1 day of age. |
| **PATHOGEN RECOGNITION** | ***TEP4*** (AGAP010812) | n-5 | i-9 | ↓ | ↑ | 32°C reduces expression beyond 1 day of age. | ↓ | ↓ | Aging-dependent decrease occurs faster at warmer temperatures. |
| **PATHWAY SIGNALING AND ANTIMICROBIAL EFFECTORS** | ***UPD3A*** (AGAP013506) | n-2 | i-1 | ↑ | ↑ | Aging-dependent increase occurs faster at warmer temperatures. | ↑ | ↑ | Aging-dependent increase occurs faster at warmer temperatures. |
| **PATHWAY SIGNALING AND ANTIMICROBIAL EFFECTORS** | ***JNK3*** (AGAP009460) | n-1 | i-2 | ↑ | ↑ | Aging-dependent increase occurs faster at warmer temperatures. | ↑ | ↑ | Warmer temperature does not increase expression at 1 day of age. |
| **PATHWAY SIGNALING AND ANTIMICROBIAL EFFECTORS** | ***NOS*** (AGAP029502) | n-2 | i-1 | – | ↑ | At the oldest age, the warmest temperature increases expression. | ↑ | ↑ | After 1 day of age, aging-dependent decrease occurs faster at warmer temperatures. |
| **PATHWAY SIGNALING AND ANTIMICROBIAL EFFECTORS** | ***DEF1*** (AGAP011294) | n-4 | i-9 | ↓ | ↑ | No interaction. | ↓ | ↓ | Aging-dependent decrease occurs faster at warmer temperatures. |
| **MELANIZATION** | ***PPO6*** (AGAP004977) | n-10 | i-9 | ↓ | ↓ | Aging-dependent decrease occurs faster at warmer temperatures. | ↓ | ↓ | Aging-dependent decrease occurs faster at warmer temperatures. |
| **MELANIZATION** | ***CLIPA5*** (AGAP011787) | n-8 | i-6 | – | ↓ | For some age groups, warmer temperature reduced expression. | – | ↓ | For some age groups, warmer temperature reduced expression. |
