## Additional File 10 for "Warmer temperature accelerates senescence by modifying the aging-dependent changes in the mosquito transcriptome, altering immunity, metabolism, and DNA repair"

**Table S3. Warmer temperature and aging interactively shape the expression of genes involved in metabolism and DNA repair.** Direction of arrow indicates upregulation, downregulation, or no change.

| | | | | NAÏVE | | | IMMUNE-INDUCED | | |
|---|---|---|---|---|---|---|---|---|---|
| | **Gene (AGAP ID)** | **Interaction K-means Cluster** Naïve | HKEC | Warmer Temp | Aging | Temperature-Age Interaction | Warmer Temp | Aging | Temperature-Age Interaction |
| **METABOLISM** | **NADH dehydrogenase (ubiquinone) Fe-S protein 8** (AGAP001711) | n-7 | i-7 | ↑ | ↓ | Warmer temperature does not increase expression at 1 day of age. | ↑ | ↓ | Warmer temperature does not increase expression at 1 day of age. |
| | **6-phosphofructokinase (*PFK*)** (AGAP007642) | n-7 | i-7 | ↑ | ↓ | The warming-based increase in expression is amplified at 15 days of age. | ↑ | ↓ | The warming-based increase in expression is amplified at 15 days of age. |
| | **Pyruvate dehydrogenase phosphatase regulatory subunit** (AGAP002217) | n-7 | i-7 | ↑ | ↓ | Warmer temperature decreased expression at 1 day of age. | ↑ | ↓ | Warmer temperature decreased expression at 1 day of age. |
| | **Isocitrate dehydrogenase (NAD+)** (AGAP002192) | n-7 | i-7 | ↑ | ↓ | Warmer temperature decreased expression at 1 day of age. | ↑ | ↓ | Warmer temperature decreased expression at 1 day of age. |
| **DNA REPAIR** | **DNA helicase MCM8** (AGAP002580) | n-6 | i-3 | ↓ | ↑ | The warming-based increase only occurs beyond 1 day of age. | ↓ | ↑ | The warming-based increase only occurs beyond 1 day of age. |
| | **DNA ligase 1** (AGAP009222) | n-5 | i-3 | ↓ | ↑ | The warming-based increase only occurs beyond 1 day of age. | ↓ | ↑ | The warming-based increase only occurs beyond 1 day of age. |
| | **DNA repair protein Rad62** (AGAP010060) | n-6 | i-4 | ↓ | ↑ | The warming-based increase only occurs beyond 1 day of age. | ↓ | ↑ | The warming-based increase only occurs beyond 1 day of age. |
| | **DNA mismatch repair protein MSH4** (AGAP012245) | n-5 | i-3 | ↓ | ↑ | The warming-based increase only occurs beyond 1 day of age. | ↓ | ↑ | The warming-based increase only occurs beyond 1 day of age. |
